# GTP biosynthesis is a therapeutic vulnerability in Rac1-mutant melanoma

**DOI:** 10.64898/2026.09.23.753957

**Authors:** Samson Eugin Simon, Tiannan Wang, Arsha Sreekumar, Zaria Yearby, Isha Patel, Varsha Tandra, Catherine C. Hedrick, Jie Li, David W. Wolff

**Affiliations:** Georgia Cancer Center, Medical College of Georgia, Augusta University, Augusta, GA; Vascular Biology Center, Medical College of Georgia, Augusta University, Augusta, GA; Immunology Center of Georgia, Medical College of Georgia, Augusta University, Augusta, GA; Department of Pathology, Medical College of Georgia, Augusta University, Augusta, GA

## Abstract

Metastatic melanoma remains highly lethal despite advances in immunotherapy and MAPK-targeted therapy. The Rac1^P29S^ mutation, present in 4-9% of cutaneous melanomas, confers intrinsic resistance to BRAF and MEK inhibitors. Like other small GTPases, Rac1 lacks suitable pockets for small molecule inhibitor binding, motivating indirect approaches to suppressing its activity. We have previously shown that wild-type Rac1 is sensitive to inhibition of *de novo* GTP biosynthesis in cancer cells, particularly suppression of the rate-limiting inosine monophosphate dehydrogenase (IMPDH) enzymes. Therefore, we asked whether IMPDH inhibition could suppress Rac1^P29S^ and its associated phenotypes in melanoma. We found that IMPDH inhibition reduced Rac1 activity, impaired Rac1-dependent phenotypes, and induced S-phase arrest in Rac1^P29S^-harboring cells. Moreover, IMPDH inhibition synergized with the BRAF inhibitor vemurafenib and the MEK inhibitor trametinib in Rac1^P29S^ melanoma cells. Dual BRAF and IMPDH inhibition also resulted in enhanced suppression of MEK and ERK phosphorylation in vitro. In syngeneic mouse models, the FDA-approved IMPDH inhibitor mycophenolate mofetil sensitized Rac1^P29S^-expressing melanoma tumors to trametinib, including tumors made refractory by prior trametinib exposure. We further identified a feed-forward circuit in which Rac1 sustains IMPDH2 expression levels through JNK and c-Jun/AP1 signaling, potentially coupling the GTPase to its own nucleotide supply. These findings establish GTP biosynthesis as a targetable vulnerability in Rac1^P29S^ melanoma and support IMPDH inhibition as a rational partner for MAPK-targeted therapy.

## INTRODUCTION

Metastatic melanoma is a highly lethal form of skin cancer. Although immune checkpoint inhibitors (ICI) have improved outcomes for those with advanced disease, a substantial fraction of patients do not respond, and most who do ultimately progress [1, 2]. Patients with progressive disease following ICI can benefit from inhibitors targeting the Mitogen-Activated Protein Kinases (MAPK) BRAF and MEK, such as vemurafenib and trametinib, respectively [3–5]. However, acquired resistance to these agents develops almost universally [6–8]. Thus, there remains a significant unmet need for improved therapeutics in patients with advanced melanoma which is refractory to MAPK blockade.

Mutations in Proline 29 of the small GTPase Rac1 are present in approximately 4-9% of cutaneous melanomas, making it the third most commonly mutated codon after *BRAF^V600^* and *NRAS^Q61^* [9, 10]. Consistent with an oncogenic role, the prominent *RAC1^P29S^* mutation is associated with more aggressive disease at presentation while conferring intrinsic resistance to BRAF and MEK inhibitors [11–14]. Rac1 cycles between an inactive GDP-bound and an active GTP-bound state, with the active form engaging downstream effectors to regulate motility, proliferation, and survival [15, 16]. The P29S mutation alters the equilibrium of Rac1 confirmations to reduce Mg^2+^ affinity, accelerate intrinsic GDP-to-GTP nucleotide exchange, and promote effector binding [17–19]. This gain-of-function mechanism is distinct from the common mutations in Ras GTPases which impair GTP hydrolysis [20]. Despite its pro-tumor function, direct pharmacological inhibition of Rac1 has proven difficult, as small GTPases lack ideal pockets for drug binding [16, 21]. This has motivated interest in indirect strategies targeting effectors downstream of Rac1^P29S^, including PAK1 and the SRF/MRTF transcriptional complex [13, 22].

An alternative approach is to target the GTP supply required for Rac1^P29S^ activation. Inosine monophosphate dehydrogenase enzymes (IMPDH1 and IMPDH2) catalyze the rate-limiting step of *de novo* GTP biosynthesis, converting IMP to XMP (Figure 1A) [23]. Thus, IMPDH enzymes are highly expressed in cancer to meet the elevated guanylate demands of proliferating cells [24–26]. Studies by us and others demonstrated that modulating GTP production through the *de novo* pathway is sufficient to suppress the activity of Rac1 and other Rho GTPases, inhibiting cancer cell invasion and tumor growth in vivo [27–31]. Importantly, we previously reported that pharmacological IMPDH inhibition reduced the activity of Rac1^P29S^ following ectopic expression in HEK293FT cells, while the hydrolysis-deficient Rac1^Q61L^ mutant was unaffected [30]. These data are consistent with a model wherein the “fast-cycling” mechanism of Rac1^P29S^ requires ongoing GTP availability to sustain its highly active state. However, whether this sensitivity extends to endogenous Rac1^P29S^ and translates to melanoma therapy remains unknown.

**Figure 1:**
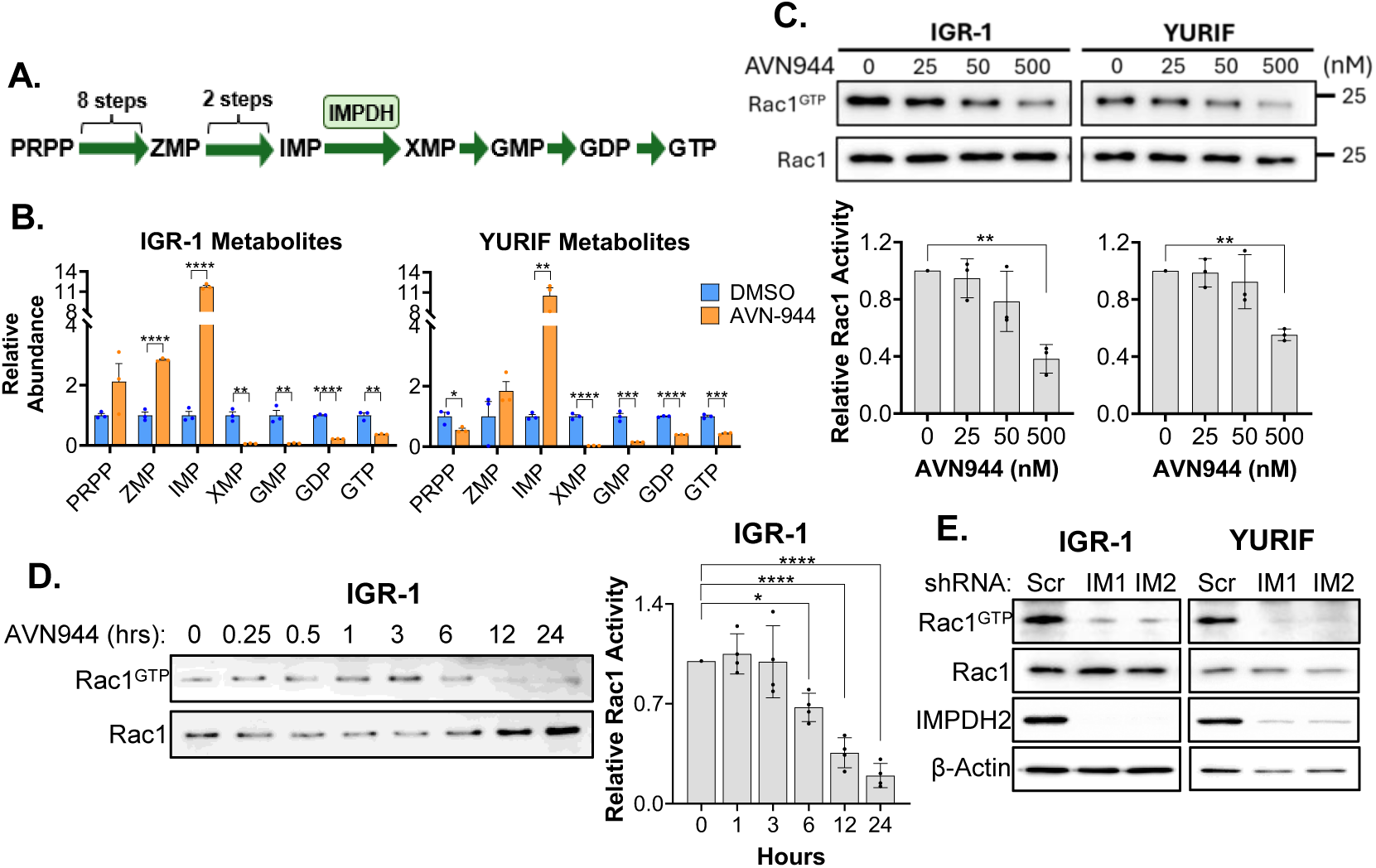
Rac1 activity in Rac1^P29S^ melanoma cells is dependent on IMPDH. **A)** Schematic depicting guanylate nucleotide biosynthesis with phosphoribosyl pyrophosphate (PRPP), 5-aminoimidazole-4-carboxamide ribonucleotide (ZMP, also called AICAR), inosine monophosphate (IMP), xanthosine monophosphate (XMP), guanosine monophosphate (GMP), GDP, and GTP shown. IMPDH enzymes catalyze the conversion of IMP to XMP. **B)** Cells were treated in biological triplicates with 500 nM AVN944 or DMSO control for 24 hours, and subjected to targeted metabolomics for the indicated metabolite. Abundance represents peak areas normalized to total protein and reported relative to DMSO control. **C)** Cells were treated with indicated concentration of AVN944 for 24 hours and probed in a Rac1 activity assay. Data is an immunoblot for both the input (total) Rac1 fraction and the bead-bound (active) fraction. (bottom) Semi-quantitative densitometry analysis of three biological replicates. **D)** Cells were treated with 500 nM AVN944 and probed in a Rac1 activity assay at the indicated timepoints, and three biological replicates were quantified as in C. **E)** Cells were transduced with lentivirus expressing two different shRNA sequences targeting *IMPDH2* (IM1, IM2) or negative (Scr) control, and probed in a Rac1 activity assay. All *p* values were determined by Welch’s t test.

Here, we report preclinical studies on IMPDH inhibition in Rac1^P29S^-positive melanoma. Utilizing Rac1^P29S^-harboring cell lines and mouse models, we observed the sensitivity of Rac1^P29S^ to reduced intracellular GTP pools. IMPDH inhibition suppressed Rac1-dependent phenotypes and increased sensitivity to MAPK-targeted therapy, including in a treatment-refractory setting. Moreover, we uncovered a feed-forward circuit in which Rac1 sustains IMPDH2 expression to reinforce its own activity in Rac1^P29S^ melanoma.

## RESULTS

### Rac1 activity depends on IMPDH-mediated GTP biosynthesis in Rac1^P29S^ melanoma

To determine whether Rac1^P29S^ activity in melanoma is sensitive to *de novo* GTP biosynthesis, we first confirmed that pharmacological IMPDH inhibition depletes cellular guanylate pools in Rac1^P29S^ harboring melanoma cells. Consistent with on-target effects of IMPDH inhibition, treatment of IGR-1 and YURIF cells with the IMPDH inhibitor AVN944 reduced the abundance of XMP, GMP, GDP, and GTP, while causing an accumulation of IMP (Figure 1A, B) [32]. We next examined whether this depletion of GTP pools translated to reduced Rac1 activity. Using standard small GTPase pulldown assays, we observed a dose-dependent decrease of active Rac1 in IGR-1 and YURIF cells following AVN944 treatment (Figure 1C). Additional experiments with a structurally distinct IMPDH inhibitor, mizoribine [33], had similar results (Supplemental Figure S1A). Moreover, we assessed the kinetics of Rac1 suppression following AVN944 treatment and found substantial reduction in Rac1 activity in IGR-1 cells after 3-6 hours (Figure 1D), with similar results in YURIF cells (Supplemental Figure S1B). Moreover, shRNA-mediated knockdown of IMPDH2 was sufficient to suppress Rac1 in IGR-1 and YURIF cells (Figure 1E). Together, these data demonstrate that Rac1^P29S^ activity in melanoma cells is highly dependent on continuous IMPDH-mediated GTP synthesis.

Active Rac1 drives invasion and migration in solid tumors. Therefore, we next asked whether inhibition of IMPDH impaired these phenotypes. In both IGR-1 and YURIF cells, treatment with AVN944 significantly reduced invasion in Boyden chamber assays (Figure 2A, Supplemental Figure S2A), and migration in wound healing assays (Figure 2B, Supplemental Figure S2B). Since macropinocytosis is governed by Rac1-dependent actin remodeling [34–36], and Rac1^P29S^ drives actin dynamics at the plasma membrane [12, 37], we further predicted that macropinocytosis would be sensitive to IMPDH inhibition in these cells. Indeed, AVN944 treatment decreased the pinocytic uptake of fluorescently labeled dextran in both cell lines (Figure 2C, Figure 2D). Together, these results demonstrate that suppression of Rac1^P29S^ activity through IMPDH inhibition is sufficient to attenuate multiple Rac1-driven phenotypes in melanoma cells.

**Figure 2:**
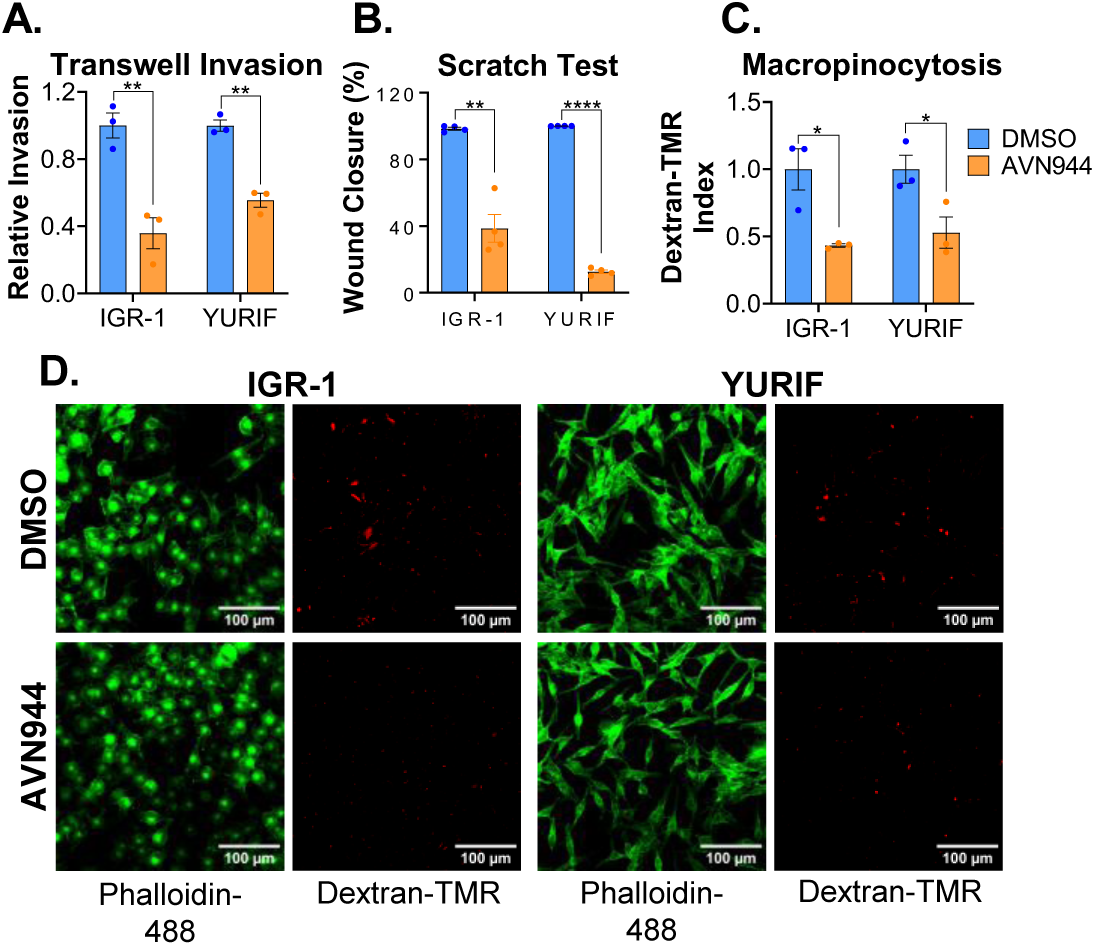
IMPDH inhibition suppresses Rac1-dependent phenotypes in Rac1^P29S^ melanoma cells. **A)** Transwell invasion assay of cells seeded into chambers in media containing 50 nM AVN944 or DMSO control. Invaded cells were normalized to methylene blue staining of cells seeded into replicate wells without chambers, and reported relative to DMSO controls. Data represents three biological replicates. **B)** Confluent cells were subjected to a wound healing assay in the presence of 50 nM AVN944 or DMSO. Data represents four biological replicates, with wound closure in the DMSO controls normalized to 100%. **C)** Cells were pre-treated with 500 nM AVN944 or DMSO control for 24 hours prior to addition of fluorescent dextran to the culture media. Pinocytic uptake of dextran was then quantified across three biological replicates with actin staining used to define cell area, and reported relative to DMSO controls. **D)** Representative micrographs of cells quantified in C. All *p* values were determined by Welch’s t test.

### IMPDH inhibition induces cell cycle arrest in Rac1^P29S^ melanoma cells

Once we established that IMPDH inhibition suppresses Rac1^P29S^ function in melanoma, we next characterized the broader consequences of AVN944 treatment in these cells. Transcriptomics data of drug-treated IGR-1 and YURIF cells revealed strong downregulation of pathways associated with proliferation, including ribosome biogenesis, chromosome segregation, and DNA replication (Figure 3A, B). Gene set enrichment analysis further confirmed a loss of proliferative gene expression in the IMPDH-inhibited cells (Figure 3C). Consistent with these observations and previous studies [32, 38–40], cell cycle analysis of AVN944-treated cells demonstrated an accumulation of cells in S-phase (Supplemental Figure S3). To determine whether these cells were actively replicating DNA, we quantified EdU incorporation on a per-cell basis. The data showed a pronounced reduction in the single cell EdU staining following AVN944 treatment in both IGR-1 and YURIF cells (Figure 3D), indicating a malfunction in DNA synthesis despite the cells accumulating in S-phase. Together, these data indicate that loss of IMPDH activity arrests Rac1^P29S^ melanoma cells in S-phase in a manner consistent with stalled replication fork progression.

**Figure 3:**
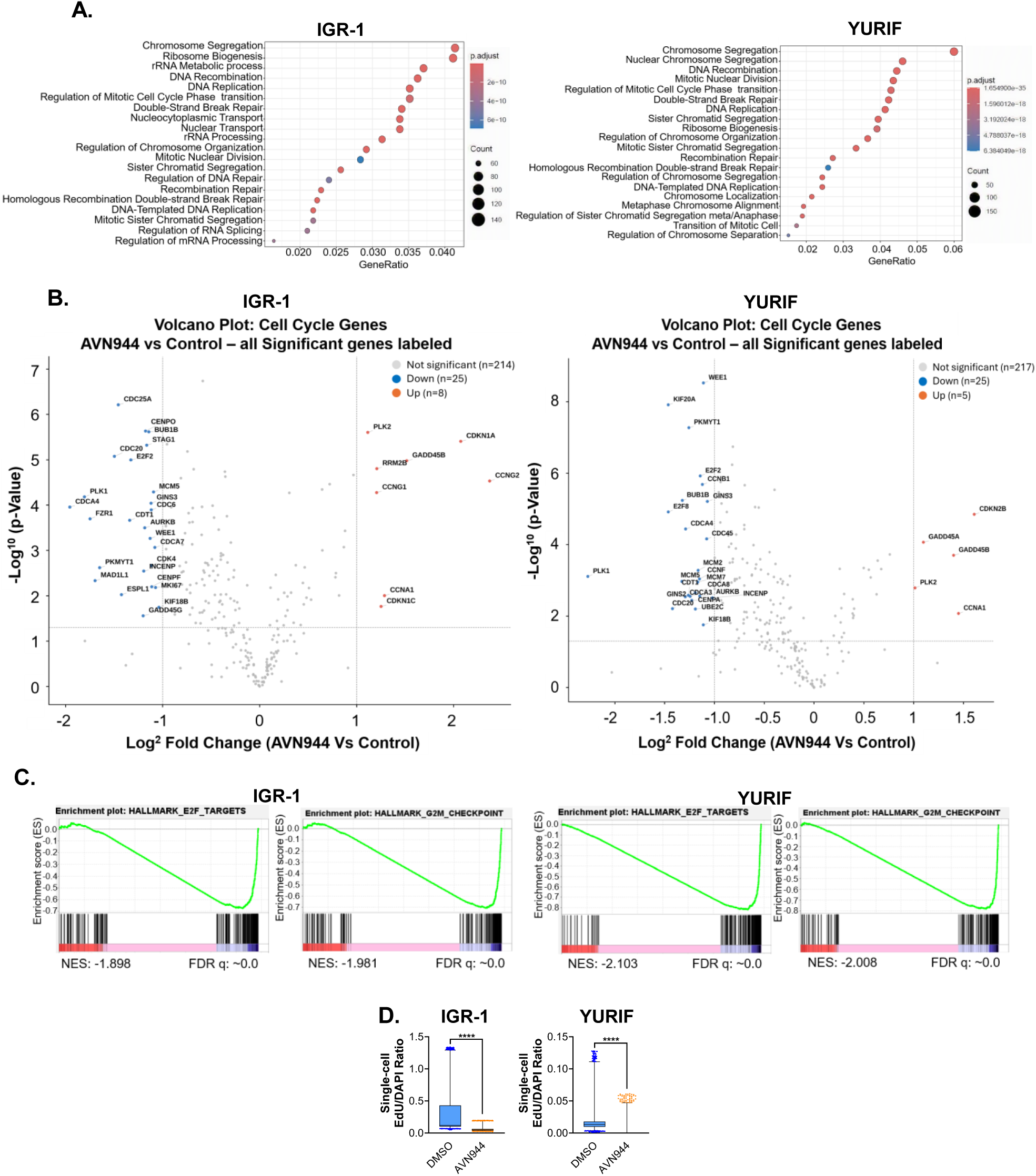
Transcriptomics reveals defective cell cycle progression in IMPDH-inhibited Rac1^P29S^ melanoma cells. **A)** Cells were treated with 500 nM AVN944 or DMSO control for 24 hours and subjected to bulk RNAseq analysis. Dotplots represent pathways enriched for suppression by AVN944. **B)** Volcano plots of genes from the MSigDB KEGG_CELL_CYCLE gene set, with genes significantly up-or down-regulated by AVN944 labeled. **C)** GSEA analysis of E2F_TARGETS and G2M_CHECKPOINT gene sets from the MSigDB Hallmark collection from AVN944-treated cells. **D)** Cells were pre-treated with 100 nM AVN944 or DMSO for 24 hours and subjected to an EdU incorporation assay. EdU intensity relative to DAPI was quantified on a per-cell basis (cell counts ranging from 4809-6431 per condition). Data is represented as a box and whiskers plot, with the box representing 25^th^-75^th^ percentiles, and the whiskers 1^st^-99^th^. Datapoints outside that range are indicated, and *p* values were determined by Mann-Whitney tests. The data presented is representative of two independent experiments.

### Targeting IMPDH synergizes with MAPK inhibition in Rac1^P29S^ melanoma cells

Since Rac1^P29S^ confers intrinsic resistance to standard-of-care BRAF and MEK inhibitors [11, 13], we asked whether IMPDH inhibition would enhance sensitivity to vemurafenib and trametinib in this setting. Thus, the *BRAF^V600E^*-harboring IGR-1 and YURIF cells were treated with increasing concentrations of AVN944, vemurafenib, or equimolar combinations of both so that total drug concentration was consistent across the three arms. MTS assays for cell viability revealed a clear leftward shift in the dose-response curves with the combination-treated cells relative to single agents, with corresponding reductions in the IC50 values across multiple biological replicates (Figure 4A). To determine whether this effect extended beyond the AVN944-vemurafenib combination, we performed similar experiments in BRAF wild-type, Rac1^P29S^-harboring YUHEF cells treated with trametinib. Indeed, the combination of AVN944 and trametinib substantially reduced the IC50 value relative to single agents (Figure 4B). Chou-Talalay analysis of the data from all cell lines indicated a synergistic effect from the combination treatments (Supplemental Figure S4A, B). These data demonstrate that IMPDH inhibition potentiates MAPK blockade in Rac1^P29S^ melanoma cells independently of BRAF status.

**Figure 4:**
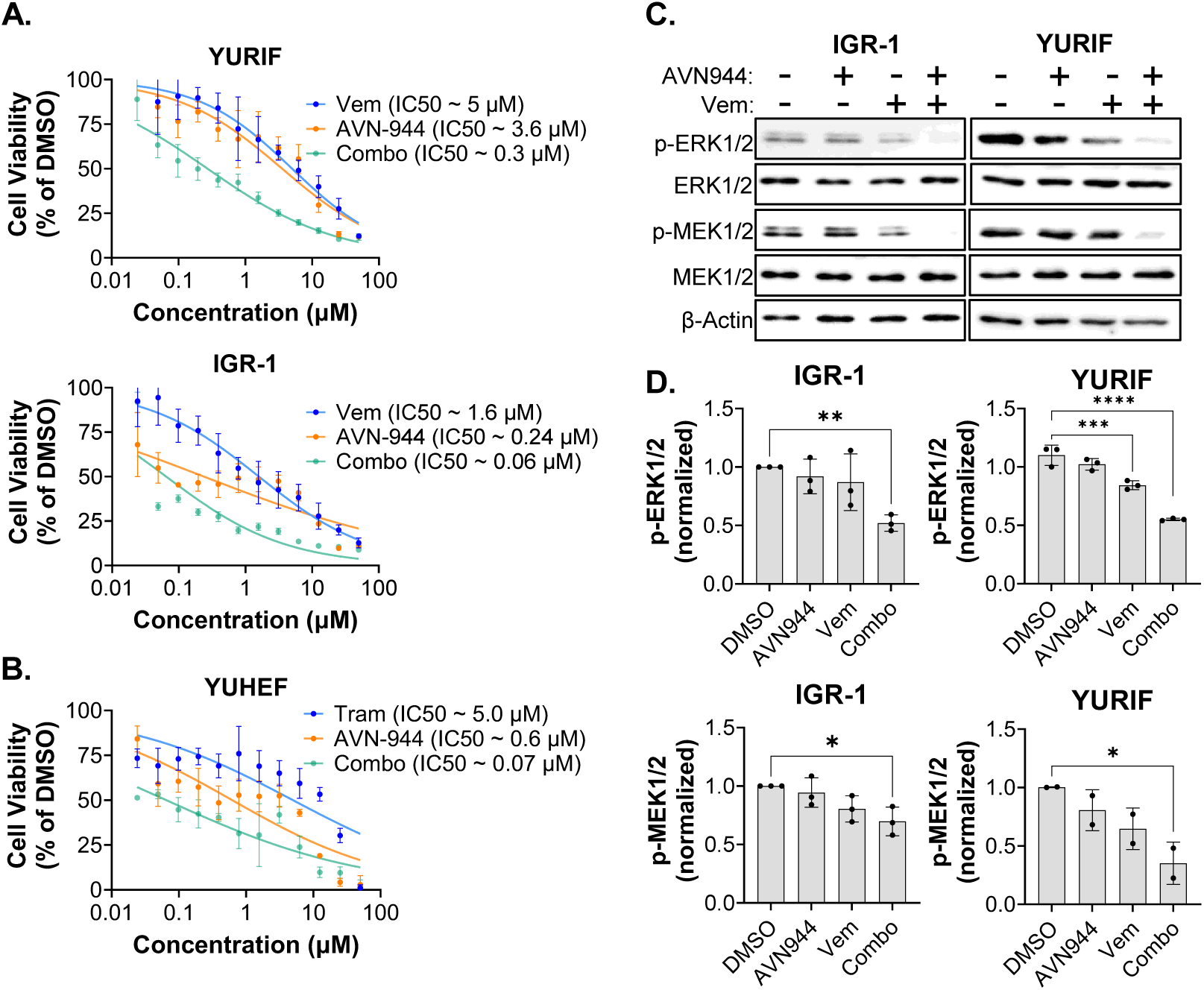
IMPDH and MAPK inhibition synergize in Rac1^P29S^ melanoma cells. **A)** Cells were treated with indicated concentrations of vemurafenib (Vem), AVN944, or the combination (combo) for 48 hours. Cell viability was determined relative to DMSO control via MTS assays (n = 6 per condition), and dose-response curves were generated via nonlinear regression analysis to calculate IC50 values. **B)** Same as A, except cells were treated with trametinib (Tram) in place of Vem. **C)** Cells were treated for 24 hours with 200 nM AVN944, 10 nM Vem, or the combination and subjected to immunoblot analyses using antibodies to the indicated proteins. β-actin was used as a loading control. **D)** Semi-quantitative densitometry analysis of biological replicates represented in C. All *p* values were determined by Welch’s t test.

We next asked whether the enhanced suppression of viability was associated with stronger inhibition of MAPK signaling. Indeed, immunoblot analysis of IGR-1 and YURIF cells treated with AVN944, vemurafenib, or the combination revealed stronger depletion of phosphorylated MEK and ERK in cells treated with the combination, observed by semi-quantitative densitometry across multiple biological replicates (Figure 4C, D). Together, these data indicate that IMPDH inhibition enhances the efficacy of MAPK-targeted therapy in Rac1^P29S^ melanoma cells, suggesting that depletion of GTP pools can partially overcome the intrinsic resistance conferred by this mutation.

### IMPDH inhibition overcomes trametinib resistance in Rac1^P29S^ melanoma in vivo

We sought to test dual IMPDH and MAPK inhibition in immunocompetent mouse models utilizing the FDA-approved IMPDH inhibitor mycophenolate mofetil (MMF), which has been in clinical use for roughly three decades as an immunosuppressant [24, 41]. Thus, we ectopically expressed Rac1^P29S^ in B16F10 cells (herein B16-P29S). Consistent with the established role of Rac1^P29S^ in MAPK inhibitor resistance, B16-P29S cells were less sensitive to trametinib than parental B16F10 cells in vitro (Figure 5A). We next established B16-P29S tumors in syngeneic C57BL/6 mice and treated them with vehicle, trametinib, MMF, or the combination. Trametinib alone had little effect on tumor growth, while MMF showed better single-agent efficacy (Figure 5B). The combination of trametinib and MMF outperformed either single agent, significantly slowing tumor growth and prolonging survival (Figure 5B, 5C). To model the clinically relevant setting of established MAPK inhibitor resistance, we next asked whether MMF could act on tumors that had already become refractory to trametinib. B16-P29S tumor-bearing mice were treated with trametinib for one week, over which tumors became refractory (Figure 5D). Refractory mice were then re-randomized to receive trametinib, MMF, or the combination. Consistent with acquired resistance, continued trametinib had no effect, and MMF alone was ineffective in this setting. In contrast, the combination of MMF and trametinib significantly slowed the growth of trametinib-refractory tumors (Figure 5D), suggesting that IMPDH inhibition can resensitize established, resistant Rac1^P29S^ melanomas to MAPK blockade.

**Figure 5:**
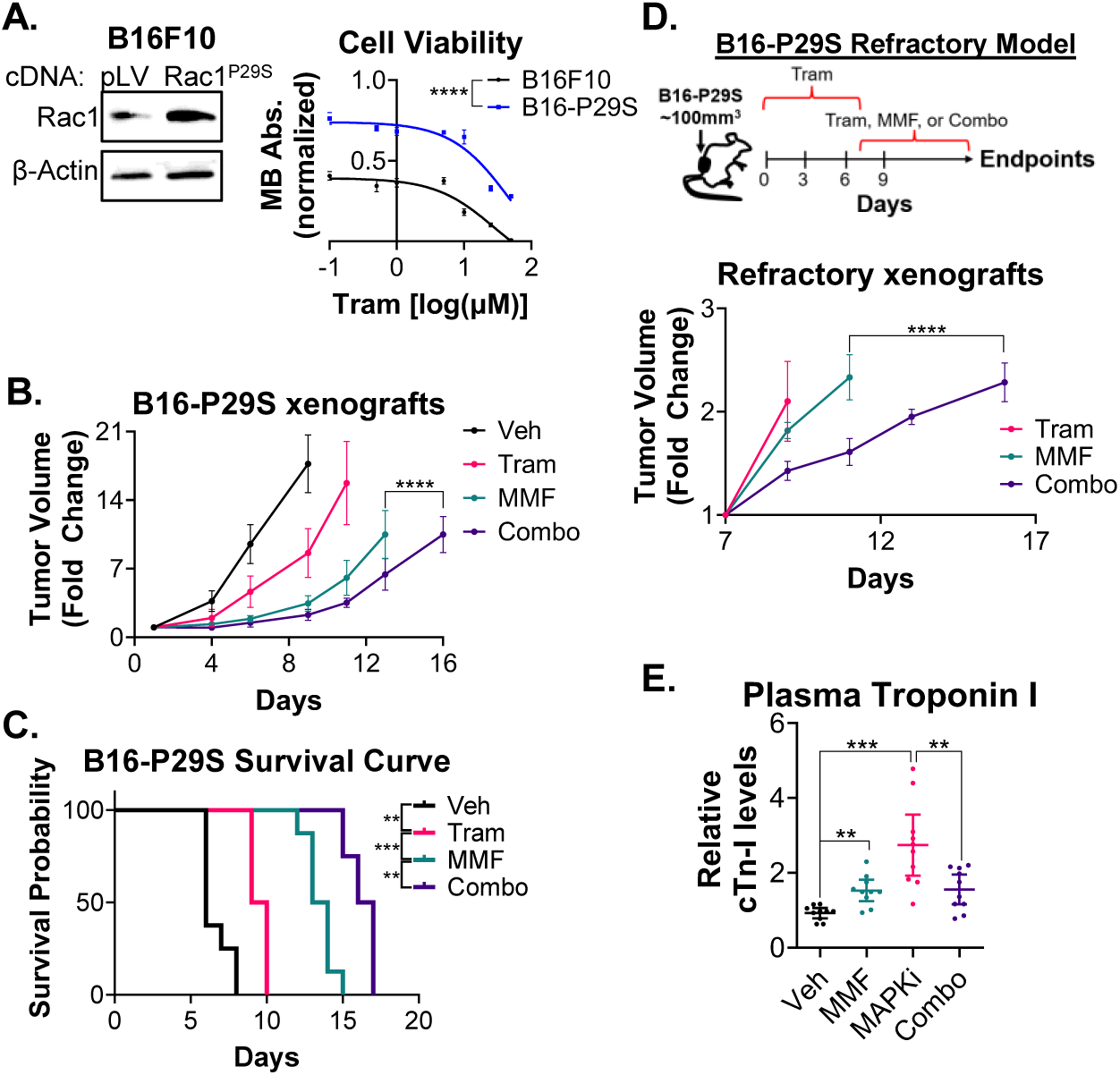
Efficacy of mycophenolate mofetil in Rac1^P29S^-expressing melanoma mouse models. **A)** B16F10 murine melanoma cells were transduced with lentivirus expressing a cDNA encoding Rac1^P29S^. (left) Immunoblot analysis confirmed ectopic expression of Rac1^P29S^. (right) Viability of parental B16F10 and Rac1^P29S^-expressing (B16-P29S) cells treated with indicated concentrations of trametinib for 48 hours. Viability was assessed via methylene blue staining, and absorbance plotted relative to DMSO controls. **B)** Growth rates of subcutaneous B16-P29S xenografts in C57BL/6 mice treated with vehicle (Veh, n =6) control, trametinib (Tram, n = 8), mycophenolate mofetil (MMF, n = 6), or the combination (Combo, n=8) over the indicated time period. Two tumors were generated and measured for each mouse. For MMF vs. combo, *p* value was determined via two-way ANOVA. **C)** Kaplan-Meier curves of mice from B pooled with a replicate experiment, with p values determined via Long-rank (Mantel-Cox) tests. **D)** C57BL/6 mice harboring B16-P29S xenografts were treated with trametinib (Tram) for 1 week and re-segregated into groups treated with Tram, MMF, or the combination (Combo). Tumor growth rate is reported with tumor volume at the time of re-segregation normalized to 1. For MMF vs. combo, *p* value was determined via two-way ANOVA. **E)** C57BL/6 mice were treated with Vehicle (Veh), trametinib + vemurafenib (MAPKi), MMF, or MAPKi + MMF (Combo) for 1-2 weeks (n = 10 per group). Blood samples were harvested from the mice and subjected to an ELISA assay for troponin I. Data is represented as a scatter plot with mean +/- 95% confidence interval, and *p* values were determined via Welch’s t tests.

Finally, we considered the cardiotoxicity associated with MAPK-targeted therapy. Combined BRAF and MEK inhibition causes cardiotoxicity in a subset of patients [42, 43]. Recent work indicates that this damage is driven by inflammation within the myocardium following MEK blockade [44]. Since MMF is a clinically established immunosuppressant with documented anti-inflammatory effects in the coronary vasculature of transplant recipients [45], we reasoned that its addition to a MAPK inhibitor regimen might mitigate cardiac injury. To test this, we treated tumor-free C57BL/6 mice with vehicle, MMF, vemurafenib plus trametinib, or the triple combination, and measured serum cardiac troponin as a marker of cardiomyocyte injury. MMF alone caused an increase in serum troponin, but combined vemurafenib and trametinib produced a substantially larger elevation (Figure 5E). Notably, the addition of MMF to vemurafenib and trametinib reduced serum troponin to levels comparable to MMF alone (Figure 5E). Thus, IMPDH inhibition may counteract inflammation-driven toxicities associated with MAPK inhibition.

### Rac1^P29S^ sustains IMPDH2 expression via c-Jun

The dependence of Rac1^P29S^ on IMPDH-derived GTP led us to ask whether Rac1 might, in turn, regulate IMPDH2. We found that shRNA-mediated knockdown of Rac1 reduced IMPDH2 protein levels in both IGR-1 and YURIF cells (Figure 6A). Next, we sought to identify the transcription factor linking Rac1 to IMPDH2 expression. Rac1 is a canonical upstream activator of the JNK-c-Jun (AP-1) signaling axis [46–48], and AP-1 consensus sites within the promoters of GTP biosynthetic enzymes have previously been proposed to contribute to their regulation [49]. Therefore, we examined the AP-1 family members c-Jun, JunB, and JunD. Knockdown of c-Jun or JunB, but not JunD, reduced IMPDH2 levels in IGR-1 and YURIF cells (Supplemental Figure S5A). We focused on c-Jun, the prototypical AP-1 subunit, for subsequent analysis, and confirmed its regulation of IMPDH2 at the protein and transcript level by immunoblot and qPCR (Figure 6B, C). Consistent with c-Jun acting downstream of Rac1, Rac1 knockdown reduced nuclear c-Jun levels in IGR-1 cells (Figure 6D). To test whether regulation of IMPDH2 by Rac1 is JNK-dependent, we ectopically expressed two different constitutively active MKK7-JNK2 fusion constructs prior to depleting Rac1 [50, 51]. Indeed, forced JNK activation with either construct rendered IMPDH2 levels insensitive to Rac1 knockdown in IGR-1 cells (Figure 6E, Supplemental Figure S5B). Together, these data suggest a feed-forward circuit in which Rac1 supports its own activity in Rac1^P29S^ melanoma via IMPDH2.

**Figure 6:**
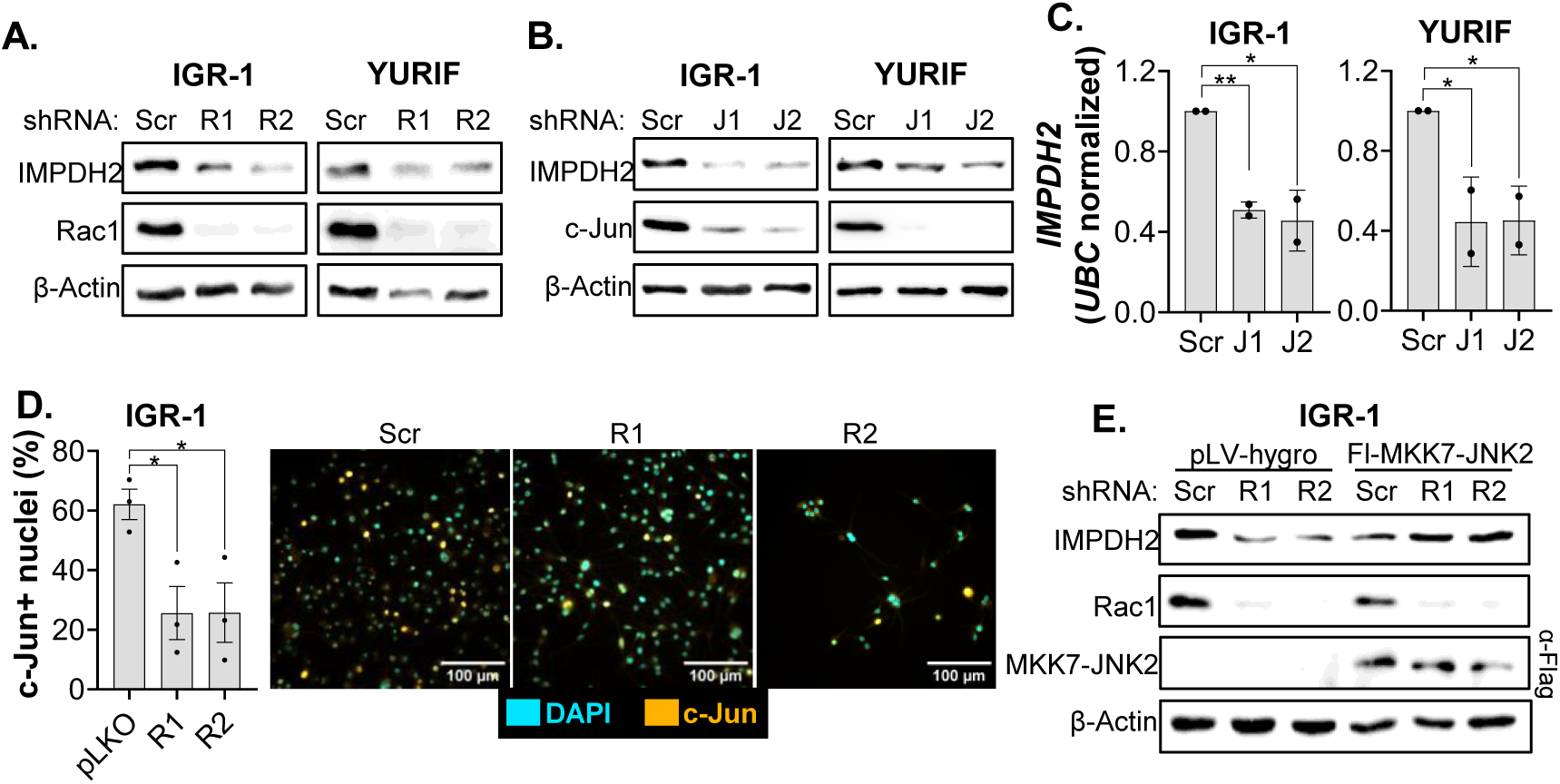
Rac1 promotes IMPDH2 expression in Rac1^P29S^ melanoma cells via c-Jun. **A)** Cells were transduced with lentivirus expressing two different shRNA sequences targeting *RAC1* (R1, R2) or negative (Scr) control, and subjected to immunoblot analysis with the indicated antibodies. β-actin was used as a loading control. **B)** Cells were transduced with lentivirus expressing two different shRNA sequences targeting *JUN* (J1, J2) or negative (Scr) control, and subjected to immunoblot analysis with the indicated antibodies. β-actin was used as a loading control. **C)** Cells were transduced as in B and assayed via qPCR with primers targeting *IMPDH2*. Primers targeting *UBC* were used as controls. Each dot represents an independent experiment measured in technical triplicates, and *p* values were determined via Nested t tests. **D)** Cells were transduced as in A and subjected to immunofluorescence analysis using anti-c-Jun antibodies with DAPI as a counterstain for nuclei. The percentage of c-Jun positive nuclei was determined across three independent experiments, and *p* values were determined via Welch’s t tests. (right) Representative micrographs of quantified cells. **E)** Cells were transduced with lentivirus expressing a Flag-tagged constitutively active JNK2 fusion protein (Fl-MKK7-JNK2) or empty vector (pLV-hygro) control. Following selection, cells were transduced a second time with lentivirus expressing shRNAs targeting *RAC1* (R1, R2) or Scr control. Cells were then subjected to immunoblot analysis with the indicated antibodies. β-actin was used as a loading control.

## DISCUSSION

Here we show that Rac1^P29S^-harboring melanoma cells depend on GTP biosynthesis to sustain Rac1 activity, and that IMPDH inhibition suppresses Rac1-driven phenotypes, arrests cells in S phase, and synergizes with MAPK-targeted therapy in vitro. In vivo, the FDA-approved IMPDH inhibitor mycophenolate mofetil (MMF) sensitized Rac1^P29S^-expressing tumors to trametinib, including in a refractory setting. We further defined a feed-forward circuit in which Rac1 supports IMPDH2 expression, potentially coupling the GTPase to its own nucleotide supply in melanoma.

We and others have previously shown that reducing *de novo* GTP synthesis through suppression of IMPDH or GMPS [27, 30, 31, 52], or by activation of GMPR [27, 28, 53], lowers the activity of Rac1 and other Rho GTPases in cancer cells. Recent work has extended this to microglia in developing mice, as postnatal administration of MMF reduced microglial branching alongside RhoA and Rac1 activity levels in the brain [54]. Moreover, we previously reported that Rac1 is sensitive to local GTP production by IMPDH2 in breast cancer cells, and proposed a model wherein Rac1 activation can be limited by GTP availability [30]. Consistent with this model, our current findings demonstrate that IMPDH inhibition suppresses Rac1 activity within several hours in Rac1^P29S^-harboring melanoma cells. While this does not rule out an indirect mechanism, the timescale is consistent with Rac1 activity becoming limited by GTP availability following IMPDH inhibition.

Our identification of a Rac1-JNK-c-Jun axis regulating IMPDH2 expression raises the question of how guanylate synthesis is transcriptionally controlled in cancer. The master regulator of cell proliferation MYC is the most well-characterized driver of *IMPDH2* transcription and nucleotide biosynthesis more broadly [49, 55–58]. Since depletion of Rac1 slows proliferation in Rac1^P29S^ melanoma cells (data not shown) [59], it is likely that a reduction in MYC activity also contributed to the decreased IMPDH2 expression we observed. Therefore, c-Jun/AP-1 and MYC may act in parallel in this setting, a possibility that warrants further study. Nonetheless, the contribution of c-Jun to IMPDH2 levels is consistent with the longstanding observation of consensus AP-1 sites within promoters of purine biosynthetic enzymes [49], and with the previously reported high activity of c-Jun in melanoma [60–62]. Further investigation into this regulation can define the precise AP-1 dimer which regulates *IMPDH2*, and in which cellular contexts the mechanism is functional. Moreover, whether the Rac1-IMPDH2 circuit is specific to Rac1^P29S^-harboring cells has not been resolved. Since the regulation we described depends on established Rac1-driven JNK-c-Jun signaling rather than a Rac1^P29S^-specific phenotype [46–48], we anticipate wild-type Rac1 can also promote *IMPDH2* expression.

The synergy we observed between IMPDH and MAPK inhibition has implications beyond Rac1-mutant melanoma. BRAF and MEK inhibitors are used across a range of malignancies such as non-small cell lung cancer, colorectal cancer, thyroid cancer, and others [63–65], but acquired therapy resistance limits their efficacy in each setting [6–8]. At the same time, most recent work on IMPDH inhibitors as potential anti-cancer agents has focused on combination treatments with genotoxic therapies such as radiation and temozolomide [66–72]. However, a growing body of evidence supports combining IMPDH inhibition with kinase-targeted therapy. Synergy between IMPDH and MEK inhibition has been reported in uveal melanoma and colorectal cancer models [73, 74], while IMPDH inhibitors synergized with EGFR blockade in non-small cell lung cancer cells [75]. Our findings extend these observations to both BRAF and MEK inhibition in Rac1^P29S^ melanoma. Importantly, the observed synergy between IMPDH inhibition and kinase-targeted therapy does not establish Rac1 suppression as the sole mechanism. The relative contribution of reduced Rac1 activity versus broader effects of GTP loss remains to be elucidated, and may vary across different malignant settings.

Notably, dual inhibition of BRAF and MEK exacerbates the toxicity profile relative to monotherapy, with adverse events occurring in essentially all patients and frequently leading to dose reduction [42, 43, 76]. In this context, recent work partly attributes cardiotoxicity to inflammation within the myocardium following MEK inhibition [44]. Consistent with the documented anti-inflammatory effects of MMF in heart transplant recipients [45], we found that its addition to a vemurafenib-trametinib cocktail attenuated the elevated plasma troponin levels caused by the MAPK inhibitors alone. This result suggests that IMPDH inhibition can attenuate a common dose-limiting toxicity of MAPK-targeted therapy, and raises the possibility that MMF could counteract other inflammation-related side effects of MAPK inhibition, such as rash and pyrexia [77, 78]. While these potential benefits must be weighed against the risk that IMPDH inhibition could compromise antitumor immune responses, the efficacy observed by us and others in immunocompetent mouse models argue that therapeutic windows exist without major detriment to antitumor immunity.

In summary, our work identifies GTP biosynthesis as a targetable vulnerability in Rac1^P29S^ melanoma and provides preclinical support for combining IMPDH inhibition with MAPK-targeted therapy. With mycophenolate mofetil FDA-approved and well tolerated, dual IMPDH and MAPK inhibition is readily testable in relevant tumor settings. More broadly, several questions arise from these findings. It will be important to define any Rac1-independent mechanisms by which IMPDH suppression synergizes with kinase-targeted therapies, testing the hypothesis that the anti-tumor potential of IMPDH inhibitors depends less on direct antiproliferative activity than on the capacity to lower GTP-dependent signaling. Moreover, while IMPDH activity supports Rac1 in a variety of contexts [27, 30, 54], it remains unknown how broadly reciprocal Rac1-IMPDH2 axis functions. Further clarifying this mechanism may inform which tumor settings are most likely to respond to IMPDH inhibitors. Finally, although IMPDH inhibitors are established immunosuppressants, evidence in mouse models suggests that IMPDH suppression does not impair responses to ICI [73, 79, 80]. The efficacy we observed in immunocompetent mice also suggests that MMF did not compromise antitumor immunity in this setting. Since ICI is standard-of-care in advanced melanoma and immunotherapy is quickly reshaping the treatment landscape across oncology, defining how IMPDH inhibition impacts antitumor immunity will be essential to further clinical development.

## MATERIALS AND METHODS

### Cell lines and cell culture

Human melanoma cell lines YURIF and YUHEF were gifted by Dr. Ruth Halaban (Yale University, New Haven, CT, USA). Human IGR-1 (catalog #300219) and murine B16F10 (catalog #305157) melanoma cells were purchased from Cytion (Sioux Falls, SD, USA). Melanoma cells were maintained in Opti-MEM^TM^ I Reduced-Serum Medium (catalog #31985070) from Thermo Fisher Scientific (Waltham, MA, USA) supplemented with 5% fetal bovine serum (FBS) and penicillin-streptomycin antibiotics. For viral packaging, Lenti-X^TM^ 293T cells (catalog #632180) were purchased from Takara Bio USA, Inc. (San Josa, CA, USA). 293T cells were maintained in DMEM supplemented with 10% FBS and penicillin-streptomycin antibiotics. All cells were maintained at 37°C and 5% CO_2_ under aseptic conditions and routinely monitored for mycoplasma contamination via MycoStrip® detection kits from InvivoGen (San Diego, CA, USA).

### Plasmids and lentiviral transduction

The PLKO.1-Scrambled control shRNA construct was a gift from Anthony Leung (Addgene plasmid #136035). For viral packaging, the pCMV-VSV-G plasmid was a gift from Bob Weinberg (Addgene plasmid #8454), and the psPAX2 plasmid was a gift from Didier Trono (Addgene plasmid #12260). The pLV-EF1a-IRES-Puro (Addgene plasmid #85132) and pLV-EF1a-IRES-Hygro (Addgene plasmid #85134) control lentiviral vectors were gifts from Tobias Meyer. The FLAG-Mkk7-JNK2-Puro vector was a gift from David Sabatini and Kris Wood (Addgene plasmid #64618). The FLAG-Mkk7-JNK2-Hygro vector was a gift from Prashant Mali (Addgene plasmid #170228). A pcDNA3.1(+) plasmid expressing Rac1^P29S^ was purchased from GenScript (Piscataway, NJ, USA), and the cDNA was cloned into pLV-EF1a-IRES-Puro utilizing the BamHI and EcoRI restriction sites. All lentiviral shRNA constructs with targeting sequences were purchased from Sigma-Aldrich (St. Louis, MO, USA) with the following clone numbers: IMPDH2 (TRCN0000026512, TRCN0000026534), RAC1 (TRCN0000318432, TRCN0000004871), JUN (TRCN0000039589, TRCN0000039590), JUNB (TRCN0000014944, TRCN0000232084), and JUND (TRCN0000416347, TRCN0000433573). Lentivirus was generated following transfection of 293T cells with psPAX2, VSV-G, and the transfer plasmid using polyethylenimine (PEI). Viral supernatants were collected 48 hours post transfection and concentrated following incubation at 4°C with polyethylene glycol. Target cells were transduced overnight in the presence of 8 µg/mL polybrene, and selected with appropriate antibiotics.

### Mouse xenograft model

All animal experiments were performed in accordance with institutional guidelines and approved by Augusta University’s Institutional Animal Care and Use Committee. C57BL/6J mice were obtained from The Jackson Laboratory (strain #000664; RRID: IMSR_JAX:000664). For xenograft experiments, 5 x 10^5^ B16F10 melanoma cells ectopically expressing Rac1^P29S^ were suspended in PBS and subcutaneously injected into the flanks of 10–12-week-old C57BL/6J mice. Tumors were allowed to reach approximately 100-150 mm^3^ before segregation into treatment groups. Mice were subsequently treated with vehicle, mycophenolate mofetil (MMF; 50 mg/kg), trametinib (Tram; 2 mg/kg), or the combination of MMF and Tram by oral gavage three times weekly. Tumors were measured via electronic caliper, and animals were monitored until tumor burden reached the predefined endpoint of 2,000 mm^3^.

### Drugs and antibodies

AVN944 (catalog #21284) and mizoribine (catalog #23128) were purchased from Cayman Chemical (Ann Harbor, MI, USA). MMF (catalog #M2387) was purchased from TCI America (Portland, OR, USA). Vemurafenib (catalog #HY-12057) and trametinib (catalog #HY-10999) were purchased from MedChemExpress (Monmouth Junction, NJ, USA).

Antibodies used in this study: anti-Rac1 (catalog #66122-1-Ig), anti-IMPDH2 (catalog #67663-1-Ig), anti-MEK1/2 (catalog #11049-1-AP), anti-DYKDDDDK (catalog #20543-1-AP), and HRP-conjugated anti-β-actin (catalog #HRP-60008) from purchased from Proteintech (Rosemont, IL, USA). Anti-c-Jun (catalog #9165), anti-ERK1/2 (catalog #9102), anti-phospho-ERK1/2 (Thr202/Tyr204) (catalog #9101), and anti-phospho-MEK1/2 (Ser217/221) (catalog #9154) were purchased from Cell Signaling Technology (Danvers, MA, USA). HRP-conjugated anti-mouse (catalog #1706516) and anti-rabbit (catalog #1706515) secondary antibodies were purchased from Bio-Rad (Hercules, CA, USA).

### Rac1-GTP pulldown assay

Active Rac1-GTP was measured using PAK1-PDB beads (catalog #PAK02) from Cytoskeleton (Denver, CO, USA). Briefly, cells were lysed in 1% NP-40 lysis buffer, and 500-1,000 μg of total protein was incubated with PAK1-PDB protein beads while rotating at 4°C according to the manufacturer’s instructions. Following incubation, beads were washed three times with cold lysis buffer to remove unbound proteins. Bead-associated and total Rac1 were analyzed via immunoblotting, and bands were visualized using the Amersham^TM^ ImageQuant^TM^ 800 western blot imaging system from Cytiva (Marlborough, MA, USA). Densitometric analysis was performed using FIJI, and Rac1-GTP signals were normalized to the appropriate total Rac1 signal.

### Mass spectrometry analysis of nucleotide pools

Melanoma cells were washed and harvested in ice-cold PBS. Then, 600 µl of ice-cold 80% methanol was added to the cell pallet together with stainless steel beads (0.9-2.0 mm diameter) and vortexed briefly. The sample was then homogenized in a Bullet Blender homogenizer at 4°C for 3 minutes, centrifuged at 16,000 x g for 15 minutes at 4°C, and 500 µl of supernatant was transferred into a new tube and vacuum dried at 30°C. The sample was reconstituted into 60 µl 20% methanol before LC-MS analysis. Separation of nucleotides was performed using an Agilent Poroshell 120 EC-C18 (100×2.1 mm, 2.7 µm) on a Shimadzu Nexera UHPLC system at a flowrate of 0.2 mL/min using a gradient elution created between buffer A (10 mM tributylamine aqueous solution adjusted pH to 4.95 with 15 mM acetic acid) and methanol, from 10% to 40% methanol in 30 minutes at room temperature. The effluent was ionized using negative ion electrospray on a TSQ Quantiva Triple Quadrupole Mass Spectrometer from Thermo Fisher Scientific (Waltham, MA, USA) with the following instrument settings: ion spray voltage 3500V, sheath gas 10, ion transfer tube temperature 350, aux gas 5, and unit resolution for Q1/Q3. The optimal collision energy and RF lens were determined using standards for each nucleotide. The integrated peak areas for the transitions were calculated for each sample using Skyline software (version 20.0, University of Washington).

### Invasion assay

Cell invasion was assessed using NEST cell culture inserts with 8.0 μm pores in polycarbonate membranes (catalog #724301) purchased from Midwest Scientific (Fenton, MO, USA). Inserts were coated in a 12 well plate with 1% Geltrex^TM^ LDEV-Free Reduced Growth Factor Basement Membrane Matrix (catalog #A1413201) diluted in serum-free DMEM at 37°C for 30 minutes. Melanoma cells (1 × 10^5^ cells per insert) were suspended in serum-free DMEM with or without IMPDH inhibitor and seeded into the upper chamber. The lower chamber contained DMEM supplemented with 10% FBS as a chemoattractant. Cells were allowed to invade through the Geltrex-coated membrane for 48 hours. Following incubation, non-invading cells were removed from the upper surface of the membrane with PBS-soaked cotton swabs. Invaded cells on the lower surface were fixed and stained with a solution of 1% methylene blue and 50% methanol. Micrographs were taken of the dried chambers, and invaded cells were quantified by assaying absorbance of the methylene blue dissolved in 1% SDS.

### Migration assay

Cell migration was evaluated using a wound healing assay. YURIF and IGR-1 melanoma cells were seeded at 5 × 10^4^ cells per well in 24-well plates and cultured for 24 h. A uniform linear wound was then generated using sterile pipette tips. Wells were washed with PBS to remove detached cells and were subsequently maintained with or without IMPDH inhibition. Images were acquired over three days. The extent of wound closure was quantified using FIJI and expressed relative to the initial wound area.

### Dextran-TMR uptake assay

Macropinocytosis activity was assessed with 70 kDa fixable dextran conjugated to tetramethylrhodamine (TMR, catalog #D1818) from Thermo Fisher Scientific (Waltham, MA, USA) using previously established protocols [81]. Melanoma cells were seeded on 8 well chambered slides and grown to approximately 70% confluency. Then, complete media was replaced with serum-free DMEM overnight with or without IMPDH inhibition. Dextran-TMR was then added to the media at a final concentration of 0.5 mg/mL for 30 minutes at 37°C and 5% CO_2_. The slides were then placed on ice, washed 5 times with ice-cold PBS, and fixed with 4% paraformaldehyde for 15 minutes at room temperature. After 3 further PBS washes, actin was stained with Phalloidin CruzFluor^TM^ 488 (catalog #sc-363791) purchased from Santa Cruz Biotechnology (Dallas, TX, USA) for 1 hour at room temperature in PBS. After additional washing, slides were mounted with coverslips and micrographs were captured on an epifluorescence microscope. In FIJI, actin images were used to define cell boundaries and create masks, and dextran-TMR uptake was quantified as a percentage of cell area positive for TMR staining.

### Cell cycle analysis

For cell-cycle analysis, treated and untreated melanoma cells were collected, washed with PBS, and passed through a 70 µm pore cell strainer to minimize cell aggregation. Cells were fixed with 70% ethanol in PBS overnight at −20°C. Following fixation, cells were collected by centrifugation and washed with PBS. Cells were then resuspended in PBS containing propidium iodide (50 μg/mL) and RNase (100 μg/mL) and incubated for 30 min at room temperature. DNA content was subsequently assayed by flow cytometry, and cell-cycle distributions were determined from DNA-content profiles using FlowJo version 10.8.1.

### EdU incorporation assay

Active DNA replication was determined by assaying incorporation of 5-Ethynyl-2’-deoxyuridine (EdU, catalog #20518) purchased from Cayman Chemical (Ann Harbor, MI, USA). Cells were seeded onto 8 well chambered slides and treated with or without IMPDH inhibitor for 24 hours, and EdU was added to the culture media at a final concentration of 10 µM for 1 hour. Then, cells were fixed with 4% paraformaldehyde for 15 minutes at room temperature, washed twice with 3% BSA in PBS, permeabilized with 0.5% Triton^TM^ X-100 in PBS for 20 minutes at room temperature, and washed again with 3% BSA in PBS. A reaction buffer was then added consisting of 50 mM Tris (pH 7.4), 150 mM NaCl, 1 mM copper (II) sulfate, 10 mM ascorbic acid, and 5 µM FAM-azide (catalog #42389) purchased from Cayman Chemical (Ann Harbor, MI, USA). Reaction proceeded protected from light for 30 minutes at room temperature. The cells were then washed once with 3% BSA in PBS, once with a solution consisting of 0.5 mM EDTA and 2 mM NaN_3_ in PBS, and once with PBS. Cells were then counter-stained with DAPI and mounted with coverslips for imaging on an epifluorescence microscope. In FIJI, the Analyze Particles function was used to quantify the green FAM and blue DAPI intensity on a per-cell basis.

### Cell viability assay

Cell viability was measured using the colorimetric CellTiter 96® Aqueous One Solution Cell Proliferation Assay (MTS) (catalog #G3582) purchased from Promega (Madison, WI, USA) according to manufacturer’s instructions. In brief, melanoma cells were seeded in 96 well tissue culture plates at 5 x 10^3^ cells per well and allowed to adhere overnight. Cells were then treated for 48 hours with compounds as indicated in the figures and figure legends. Following treatment, 20 μL of MTS reagent was added to each well containing 100 μL of medium and incubated at 37°C for 2 hours. Absorbance was measured at 490 nm using a microplate reader. Viability was expressed relative to the appropriate vehicle-treated control.

### RNA extraction and gene expression analysis

Total RNA from melanoma cells were extracted using the GeneJET RNA Purification Kit (catalog #K0731) purchased from Thermo Fisher Scientific (Waltham, MA, USA). Then, 2 μg of purified RNA was reverse transcribed into complementary DNA (cDNA) using the High-Capacity cDNA Reverse Transcription Kit by Applied Biosystems™ (catalog #4368814) purchased from Thermo Fisher Scientific (Waltham, MA, USA). To assay gene expression, qPCR was performed with SYBR green master mix on a CFX Opus 96 Real-Time PCR System from Bio-Rad (Hercules, CA, USA). Quantification was performed using the ΔΔCt method using *UBC* as the reference gene. The primer sequences for qPCR used in this study (forward and reverse) were as follows: *IMPDH2* (CATGGCCGACTACCTGATTAG, CAGTGAAGTCGATGTACCCAG), and *UBC* (ACCAGCAGAGGCTGATCTTT, TGATGGTCTTGCCAGTGAGT).

### Bulk RNAseq analysis

RNA isolation, library preparation, sequencing, and standard bioinformatics analyses were performed by Admera Health (South Plainfield, NJ, USA). Experimental samples were pelleted in Monarch® DNA/RNA protection reagent (catalog #T2010) purchased from New England Biolabs (Ipswich, MA, USA) and snap-frozen prior to shipping on dry ice. After RNA isolation and QC, sequencing libraries were prepared using the NEBNext® Ultra^TM^ II Directional RNA Library Prep Kit for Illumina® from New England Biolabs (Ipswich, MA, USA) coupled with Poly A selection. Prepared libraries were sequenced on an Illumina NovaSeq platform (Illumina, San Diego, CA, USA) utilizing a 2 × 150 bp paired-end (PE) configuration, with a sequencing depth of approximately 40 million total PE reads per sample. After raw sequencing data (FASTQ files) underwent FastQC (version v0.11.8), reads were mapped to the human reference genome (GRCh38/hg38) using STAR aligner (version 2.7.1a). Picard tools (version 2.20.4) was applied to mark duplicates, and StringTie (2.0.4) was used to assemble RNA-seq alignments into potential transcripts. FeatureCounts (version 1.6.0)/HTSeq was used to count mapped reads for genomic features such as genes, exons, promoters, gene bodies, genomic bins and chromosomal locations. De-Seq2 (version 1.14.1) was used to do differential expression analysis. Gene Ontology Analysis was done using ClusterProfiler package in R [82]. Volcano plots were generated in Python (v3, pandas v3.0.2, SciPy v1.17.1, Matplotlib v3.10.8), plotting log_2_ fold change against −log_10_(p-value) for all matched cell cycle genes identified using the KEGG “Cell cycle” pathway (hsa04110), retrieved via the MSigDB KEGG_CELL_CYCLE gene set [83, 84], and matched to the previously normalized expression dataset by gene symbol. The significantly up-and down-regulated genes were highlighted and labeled based on a nominal significance threshold of p < 0.05 and an absolute log_2_ fold change ≥1. GSEA v4.3.2 (Broad Institute) analysis was performed on pre-ranked gene lists (treatment vs. control) using the MSigDB Hallmark gene set collection (Homo sapiens, v2026.1.Hs). Genes were ranked by Signal2Noise metric, and enrichment significance was assessed using 1,000 gene-set permutations, with enrichment scores normalized for gene set size (NES).

### Cardiac troponin I measurement

Blood samples were collected from 12-14-week-old female C57BL/6 mice (n = 10 per group, with data pooled across two independent experiments) after treatment with the indicated drugs for 7-14 days. Samples were collected into EDTA-containing tubes and centrifuged at 3,000 × g for 15 min to separate plasma. Cardiac troponin I (TNNI3) concentrations were measured using the Mouse Cardiac Troponin I ELISA Kit (catalog #EEL112) purchased from Thermo Fischer Scientific (Waltham, MA, USA). According to the manufacturer’s instructions, plasma samples were normalized to equal protein concentrations and diluted to 1,000 pg/mL with the assay dilution reagent. Diluted plasma samples and cardiac troponin I standards were added to antibody-coated wells and incubated as specified. The wells were washed, incubated with the enzyme-linked detection reagent, and treated with substrate before the reaction was stopped. Absorbance was measured at 450 nm using a microplate reader. Cardiac troponin I concentrations were calculated from the standard curve according to the manufacturer’s instructions.

### Quantification and statistical analysis

Statistical analyses were performed using tools within GraphPad Prism. Statistical tests used for individual experiments are indicated in the corresponding figure legends. A two-tailed *p* value of <0.05 was considered statistically significant, with * denoting *p* < 0.05, ** denoting *p* < 0.01, *** denoting *p* < 0.001, and **** denoting *p* < 0.0001. Unless otherwise indicated, graphs represent mean +/- standard deviation from multiple biological replicates.

## ACKNOWLEDGMENTS

This work was supported Georgia Cancer Center start-up funds and grants from the National Institutes of Health to D.W.W. (NCI R00CA266920 and sub-award of NHLBI U54HL169191). Z.Y. was supported by the Summer Scholar Program through the Center for Undergraduate Research and Scholarship at Augusta University. J.L. was supported by grants from the National Institutes of Health (NHLBI R01HL146807) and the American Heart Association (23TPA1077767). V.T. was supported by a postdoctoral fellowship from the American Heart Association (24POST1196232). The authors thank the Georgia Cancer Center Flow and Mass Cytometry Core Facility (RRID:SCR_025747) for technical support in determining cell cycle distribution, and the Augusta University Proteomics and Mass Spectrometry Core Facility (RRID:SCR_027673) for development of the panel for nucleotide metabolomics.

## AUTHOR CONTRIBUTIONS

S.E.S., T.W., A.S., Z.Y., I.P., V.T., and D.W.W. performed experiments. V.T. and J.L. contributed technical and subject-matter expertise to the murine troponin experiments. C.C.H. helped develop the B16-P29S model. S.E.S., T.W., A.S., and D.W.W. analyzed the data. D.W.W. conceived and oversaw the study. S.E.S. and D.W.W. wrote the manuscript with input from all authors.

## SUPPLEMENTAL FIGURE LEGENDS

**Supplemental Figure S1:** IMPDH inhibitors suppress Rac1 activity in Rac1^P29S^ melanoma cells. **A)** Cells were treated with indicated concentration (in µM) of AVN944 or mizoribine (MZN) for 24 hours and probed in a Rac1 activity assay. (left) Representative result from three independent experiments. (right) Semi-quantitative densitometry analysis of the three biological replicates, with *p* values were determined via Welch’s t tests. **B)** Cells were treated with 500 nM AVN944 and probed in a Rac1 activity assay at the indicated timepoints, and two biological replicates were quantified as in A.

**Supplemental Figure S2:** Suppression of invasion and migration in Rac1^P29S^ melanoma cells following IMPDH inhibition (related to Figure 2). **A)** Representative micrographs of stained transwells quantified in Figure 2A. **B)** Representative micrographs of scratch tests quantified in Figure 2B.

**Supplemental Figure S3:** AVN944 induces S-phase arrest in Rac1^P29S^ melanoma cells. Cells were treated with indicated concentrations of AVN944 for 24 hours and subjected to flow cytometry analysis following fixation and staining with propidium iodide. The portion of cells in different cell cycle phases was quantified from the resulting histogram.

**Supplemental Figure S4:** Synergy between IMPDH and MAPK inhibition in Rac1^P29S^ melanoma cells (related to Figure 4). **A, B)** Median-effect plots and combination index values derived from cell viability data depicted in Figures 4A and 4B.

**Supplemental Figure S5:** c-Jun promotes IMPDH2 expression levels. **A)** Cells were transduced with lentivirus expressing two different shRNA sequences targeting *JUN, JUNB, JUND,* or negative (PLKO) control, and subjected to immunoblot analysis with IMPDH2 antibodies. β-actin was used as a loading control. B) Cells were transduced with lentivirus expressing a Flag-tagged constitutively active JNK2 fusion protein (Fl-MKK7-JNK2) or empty vector (pLV-puro) control. Following selection, cells were transduced a second time with lentivirus expressing shRNAs targeting *RAC1* (R1, R2) or Scr control. Cells were then subjected to immunoblot analysis with the indicated antibodies. β-actin was used as a loading control.

