## Supplemental Figures for "GTP biosynthesis is a therapeutic vulnerability in Rac1-mutant melanoma"

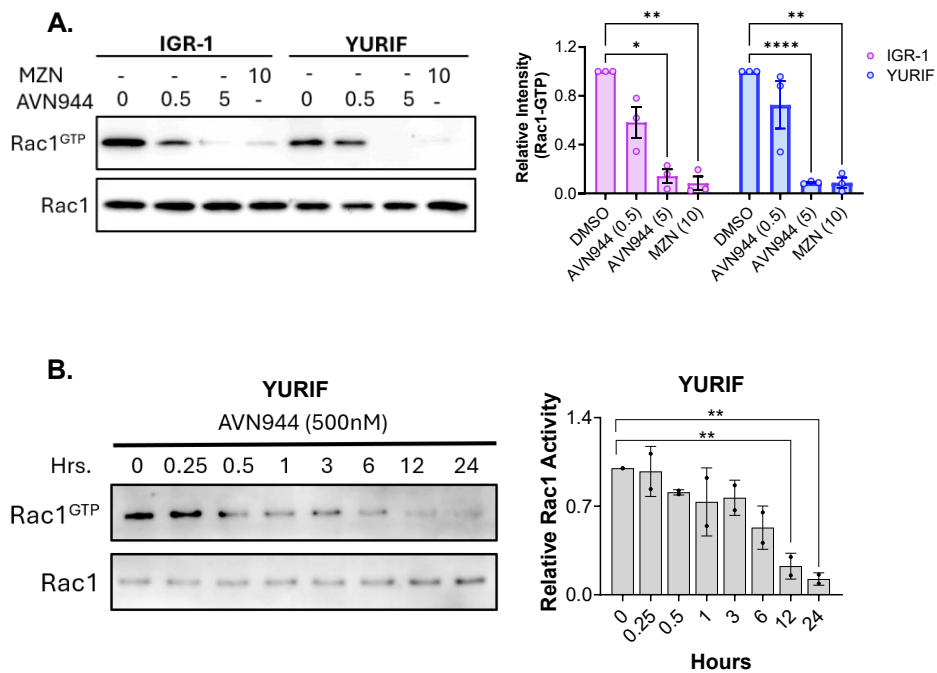

Supplemental Figure S1

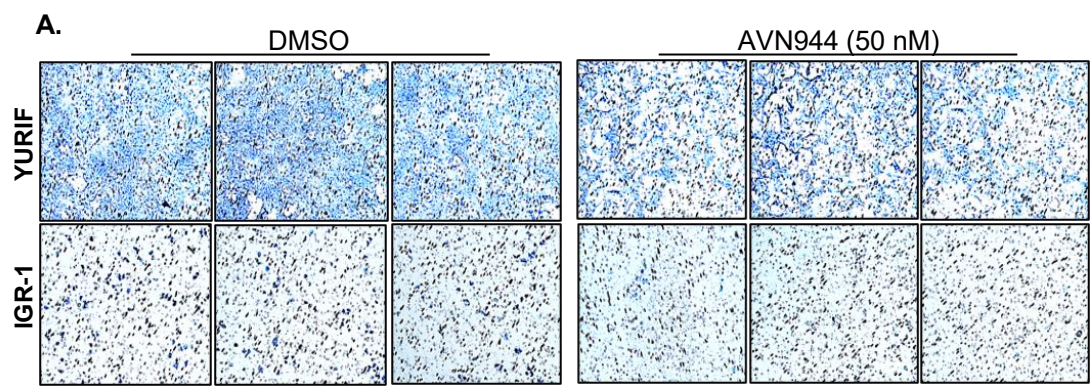

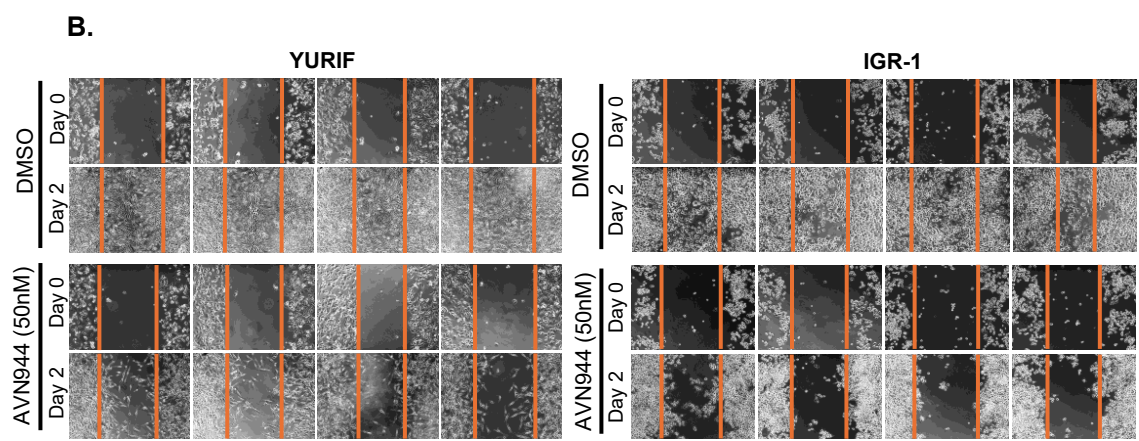

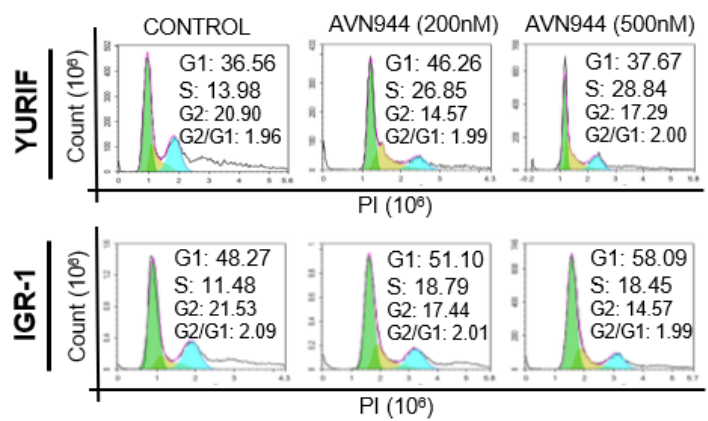

Supplemental Figure S3

**A.**

**IGR-1: Median-Effect Plot**

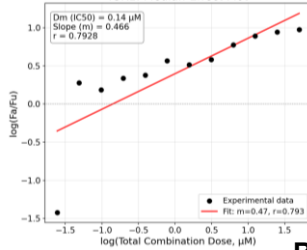

**IGR-1 Synergy**

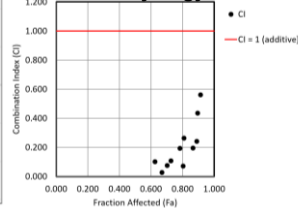

**YURIF: Median-Effect Plot**

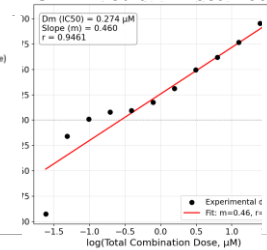

**YURIF Synergy**

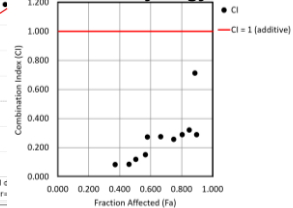

**B.**

**YUHEF: Median-Effect Plot**

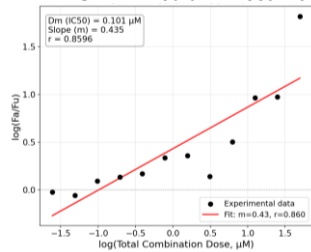

**YUHEF Synergy**

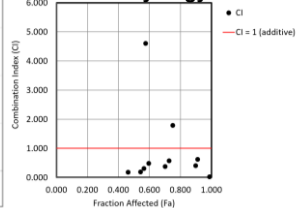

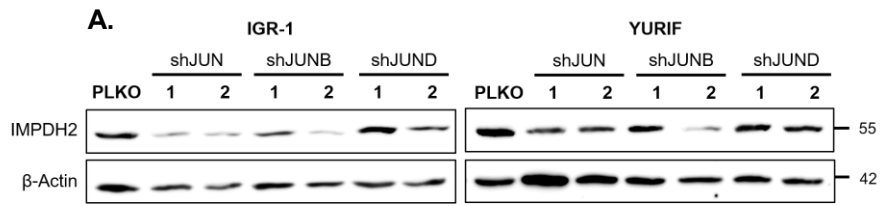

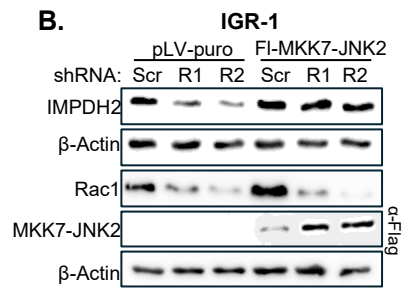
